# Distributed and drifting signals for working memory load in human cortex

**DOI:** 10.1101/2025.09.15.676305

**Authors:** Kirsten C.S. Adam, Edward Awh, John T. Serences

## Abstract

Increasing working memory (WM) load incurs behavioral costs, and whether the neural constraints on behavioral costs are localized (i.e., emanating from the intraparietal sulcus) or distributed across cortex remains an active area of debate. In a pre-registered fMRI experiment, 12 humans (12 scanner-hours each) performed a visual WM task with varying memory load (0-4 items). We replicated a localized, load-dependent increase in univariate BOLD activity in parietal cortex. However, we also observed both systematic increases and decreases in univariate activity with load across the visual hierarchy. Importantly, multivariate activation patterns encoded WM load regardless of the direction of the univariate effect, arguing against a restricted locus of load signals in parietal cortex. Finally, we observed representational drift in activity patterns encoding memory load across scanning sessions. Our results suggest a distributed code for memory load that may be continually refined over time to support more efficient information storage.

## Introduction

Working Memory (WM) is the ability to temporarily hold information in mind to guide behavior, and can only sustain high-fidelity representations of 3-4 items (Adam et al., 2017; Cowan, 2001; Luck & Vogel, 1997). This limited capacity places constraints on everyday cognitive operations like visual search, but also on more complex operations associated with general intelligence (Conway et al., 2003; Unsworth et al., 2014). Given the central role of WM capacity limits in human cognition, there is a longstanding interest in understanding the neural processing constraints that give rise to behavioral load effects.

Theories of the neural basis of working memory behaviors recognize that a distributed set of cortical regions contribute to WM. For example, early PET and fMRI work revealed activation in a broad network of areas with working memory demands, particularly the frontal cortex and a broader ‘multiple demands’ network (Duncan & Owen, 2000; Wager & Smith, 2003). However, when it comes to explaining WM *load effects*, parietal cortex has played an outsized role. Early work using functional magnetic resonance imaging (fMRI) supported a unique role for the intraparietal sulcus (IPS) in generating visual working memory capacity limits (Linden et al., 2003; Mitchell & Cusack, 2008; Todd & Marois, 2004, 2005; Xu & Chun, 2006). Increases to univariate activity in the IPS track working memory load (Todd & Marois, 2004; Xu & Chun, 2006), and changes to univariate activity in IPS also selectively predict individual differences in WM capacity limits (Todd & Marois, 2005). Single-unit recordings from frontal and parietal cortices in non-human primates have similarly found that processing constraints within parietal cortex may contribute to behavioral capacity limits (Buschman et al., 2011). Together, this body of work has led to the popular account that the parietal cortex, more so than other cortical regions, contributes to behavioral WM capacity limits and should be uniquely impacted by working memory load manipulations.

In contrast to early univariate studies, later multivariate fMRI studies of WM cast doubt on the notion that the parietal cortex plays a privileged role in facilitating working memory storage. Instead, multivariate studies of WM have found that the identity of an item held in WM can be decoded in a distributed set of cortical regions (Christophel et al., 2017), and evidence from these studies has led to sensory recruitment theories, which pose that early sensory regions are recruited, in conjunction with top-down control signals from parietal and frontal cortex, to support WM maintenance (for review, see: Adam et al., 2022). Work in this vein has consistently found that a remembered orientation or location can be decoded in cortical areas from V1 through IPS and frontal cortex (Ester et al., 2015; Harrison & Tong, 2009; Serences et al., 2009; Sprague et al., 2014, 2016), even in the absence of significant univariate activation (Serences et al., 2009). A key limitation of extant work, however, is that multivariate studies have focused on decoding only 1 or 2 items held in mind, making it diflcult to assess how working memory load impacts early sensory regions. Given the presence of single-item decoding across the visual hierarchy, an alternate account is that working memory load impacts many stages of the processing hierarchy.

In the current study, we examined the effect of working memory load on univariate and multivariate signatures of working memory in retinotopically organized regions of cortex (V1 to IPS1-4). In pre-registered analyses, we examined (1) univariate signals as a function of working memory load, while controlling for potential visual confounds via a retro-cue design, and (2) whether information about the load condition can be decoded from different areas of visual cortex, as has recently been suggested by large-scale EEG signatures of load (Adam et al., 2020; Thyer et al., 2022). In addition to testing pre-registered hypotheses about WM load signals, we (1) conducted simulations to develop a potential explanation for seemingly divergent patterns of results between univariate and multivariate metrics across the visual hierarchy, and (2) we leveraged our large dataset to test for the putative presence of representational drift in activations patterns associated with WM load. Recent work has highlighted that neural representations related to perception, decisions, memories, and actions undergo dynamic reconfiguration over time, even after measurable changes to behavioral performance have stabilized (Deitch et al., 2021; Driscoll et al., 2017, 2022; Kentros et al., 2004; Micou & O’Leary, 2023; Roth & Merriam, 2023; Schoonover et al., 2021). Working memory signals may similarly undergo dynamic changes that may be observed with human fMRI. This possibility remains under-explored in the human neuroimaging literature, in part because of the rarity of fMRI datasets with a sufficient number of experimental sessions.

To preview results, we found univariate and multivariate signatures of working memory load that are consistent with a pervasive influence of WM capacity constraints on WM representations across the visual stream. Although early fMRI work emphasized the potentially unique role of IPS in generating working memory capacity constraints on the basis of univariate increases in behavior, here we demonstrate that modulation of neural activity by working memory load is evident across the visual hierarchy. Further, we demonstrate that increases to univariate activity do not represent a reliable index of whether information about WM load is present in patterns of multivariate activity measured from each region. Rather, different regions may yield opposing univariate effects despite containing reliable multivariate information about increasing WM load. To unpack the opposing univariate and multivariate results we obtained, we used simulations to demonstrate how altering minor parameters of modelled population activity may lead to opposite univariate effects (i.e., net increase vs. decrease) without changing core features of the model (e.g., a normalization model with an activity peak for each remembered item). Finally, we found that multivariate signatures of working memory load showed directional drift as a function of the spacing between scanning sessions (spanning days to weeks), providing an intriguing departure point for future examinations of putative signatures of representational drift using large-scale population codes measured with fMRI.

## Results

Participants completed a working memory task while fMRI data were collected. Critically, we used a retro-cue task design that allowed us to examine load-related signals in cortex while controlling for the number of initially encoded items (Figure 1). On each trial, participants encoded four colored squares into working memory, and then received a retro-cue indicating which items should be stored and which could be dropped from memory.

**Figure 1.**
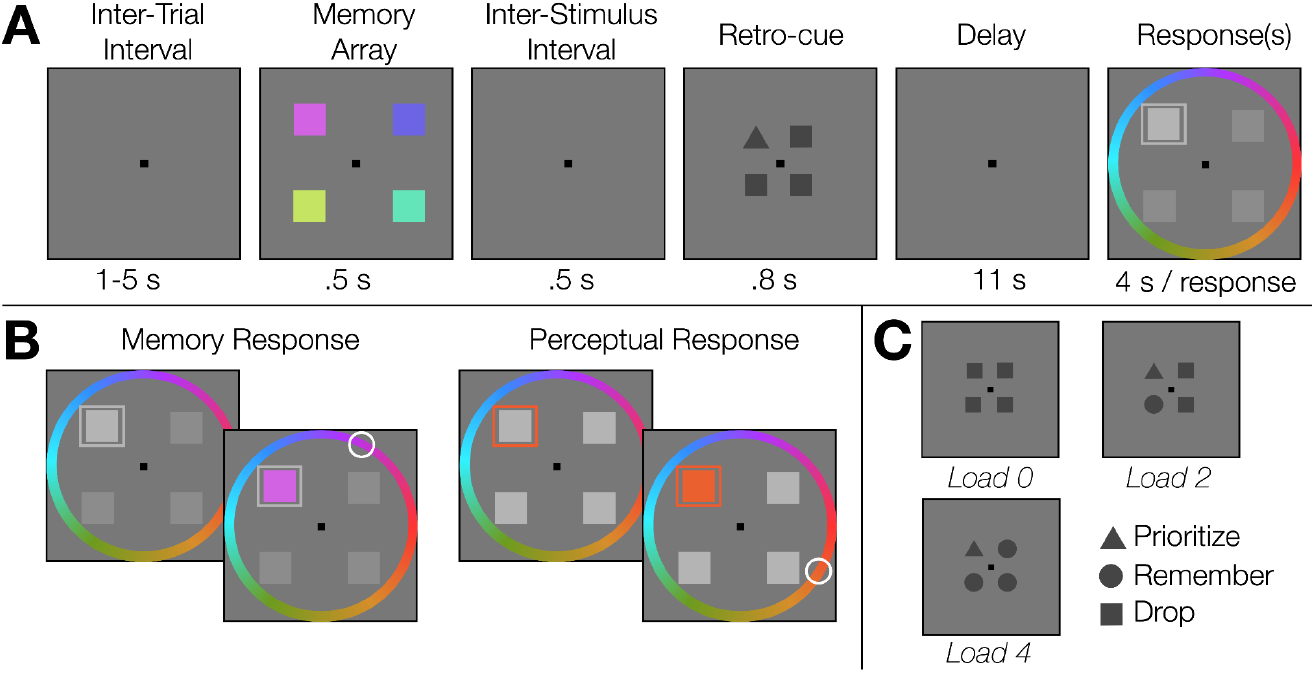
Illustration of the main working memory task. (A) Sequence of trial events for an example Load 1 trial. Participants encoded 4 colored squares into working memory, and then were instructed via retro-cue which items to store versus drop. After a delay, participants reported the remembered color(s) using a color-wheel. (B) Examples of memory responses and perceptual responses. For set sizes 1, 2 and 4, we collected 1, 2, and 4 memory responses, respectively. For set size 0 and set size 1, we collected 1 perceptual matching response, where participants matched the color of a box surrounding a randomly chosen location. Participants used a gamepad to report which color belonged at each of the remembered locations. (C) Example retro-cue arrangements for the other load conditions (Load 0, 2 and 4). The retro-cue shape indicated whether participants should drop the item (“drop”) or remember it (“prioritize”, “remember”). The cue shapes were counterbalanced across participants.

### Replication of standard behavioral load effects

We found expected working memory load effects, indicating that participants successfully used the retro-cue to selectively store items. Working memory performance decreased as working memory load increased, *F*(1.52,16.68)^1^ = 28.9, *p* < .001, *η^2^ _p_* = .72, in a linear and monotonic fashion (p<.001), and all set sizes were different from one another (p_holm_ < .03). To test how well participants used the prioritization cue, we ran a repeated measures ANOVA with factors set size (2 or 4) and prioritization (prioritized item vs other items). The prioritized item was remembered significantly better than the other items, *p =* .022. In addition, there was an interaction of set size and prioritization (*p* = .009), where the prioritization effect was larger for set size 4 than 2. Finally, we found that memory performance decreased as a function of response order for set size 4 (p < .001) but not for set size 2 (p = .23).

### Univariate load signals beyond parietal cortex

We observed heterogeneous effects of load on univariate activity across regions, as indicated by a significant 2-way interaction between Region and Load, *F*(18,198) = 39.4, *p* < .001, *η^2^ _p_* = .78. As expected, we observed a robust increase in sustained delay period activity in IPS1-4 as a function of working memory load, F(3,33) = 89, *p* < .001, *η^2^ _p_* = .89). However, load-related changes to univariate activity were not isolated to parietal cortex. Most notably, we observed robust load-related *decreases* to univariate activity in V3 (*p*<.001), V3AB (*p*<.001^1^), and hV4 (*p*=.004). We found no effect of working memory load on univariate activity in V1 (*p*=.06^1^), V2 (*p*=.50), or IPS0 (*p*=.08^1^). Figure 2A shows the univariate activity for each load condition during the delay period of the main working memory task, and Figure 2B shows baselined memory-related activity (Load 1-4 minus Load 0).

**Figure 2.**
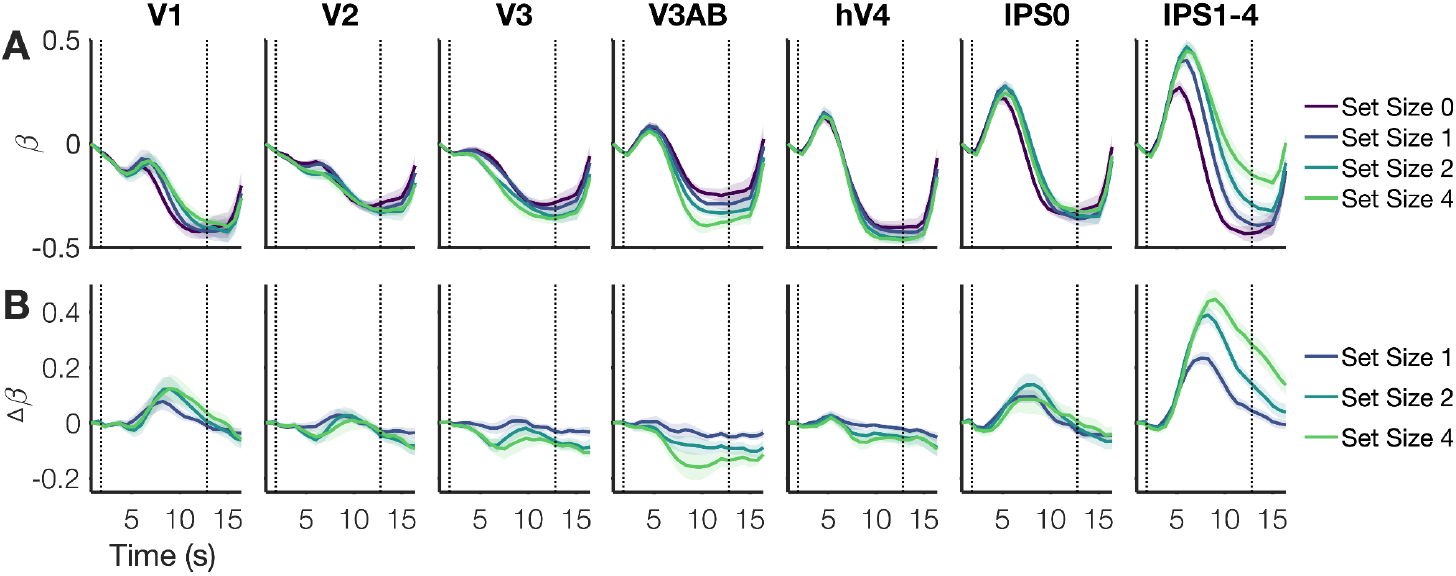
Univariate activity as a function of memory load. (A) Univariate activity (beta coefficients) for each ROI and load condition. Shaded error bars represent +/-1 SEM. Time 0 represents the onset of the memory array; dotted vertical lines indicate the beginning and end of the delay period. (B) Difference plots for each memory condition using Set Size 0 as a baseline.

### Multivariate decoding of working memory load

Although we observed positive, negative, and null effects of working memory load on univariate activity across retinotopic regions, in all regions we could reliably decode working memory load using the multivariate pattern of activity across voxels during the delay period (Figure 3A, *t*(11) > 12.4, *p* < .001, Cohen’s d > 3.59). Load decoding was not driven by an “all or none” engagement of working memory (i.e., load 0 versus all other load conditions). Rather, pairwise classification was significant for all possible pairs of load conditions (*p* < .001), and the discriminability of load condition pairs monotonically increased as the distance between load conditions increased, e.g., a difference of 1 item (0 vs. 1; 1 vs. 2) versus 2, 3, or 4 items (Figure 3B, *F*(1.1, 11.8) = 59.64, *p*<.001, *η^2^ _p_* = .84)^2^. Finally, load decoding was sustained throughout the delay period in all ROIs (Figure 3C, *p* < .001, Bonferroni-corrected comparisons shown at each timepoint). Cross-temporal generalization of decoding is shown in Figure S1.

**Figure 3.**
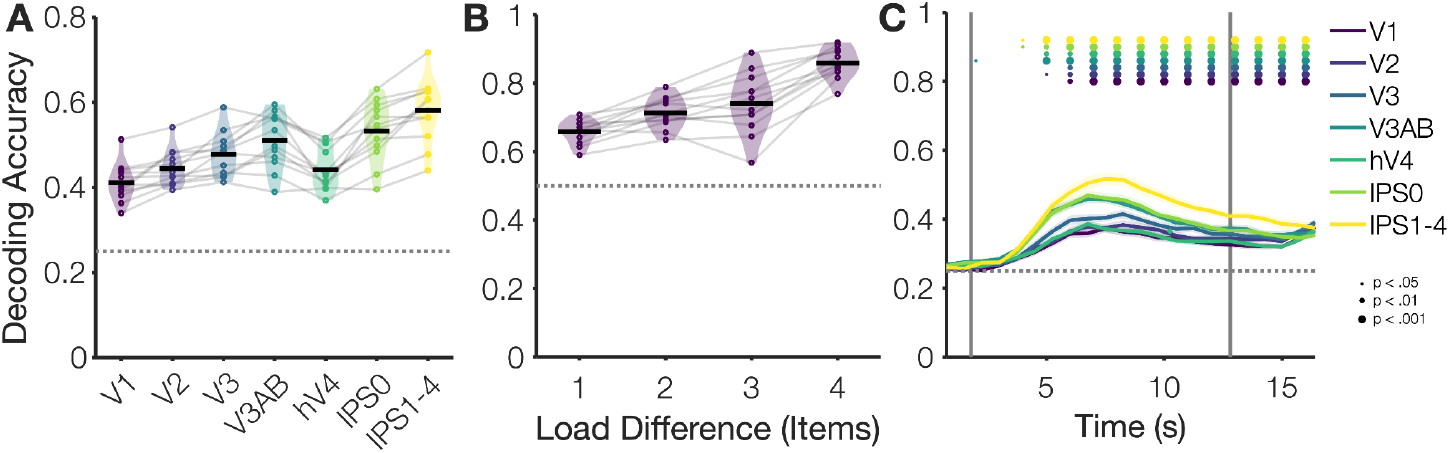
Decoding working memory load condition. (A) Average delay period decoding for each region of interest (5.25 - 12.75 s). Chance level is 25% (four load conditions). Thin gray lines represent individual subjects; violins show the distribution density. (B) Average delay period decoding for load condition pair (5.25 - 12.75 s; Chance = 50%). (C) Decoding over time for each region of interest. Chance level is 25% for decoding between the four load conditions (0, 1, 2, and 4). Decoding was performed on data averaged for a sliding time-window of 4 TRs (3 s).

### Simulating the effects of normalization and tuning properties on population responses

So far, we have found that the univariate signals showed heterogeneous effects of load, with some areas showing increase in activity, some showing decreases in activity, and some showing null effects. Despite this heterogeneity, we were able to decode the working memory load condition throughout the delay period. The ability to broadly decode working memory load is consistent with much prior work showing that the identity of a single remembered item can be decoded broadly across retinotopically organized cortex (e.g., Sprague et al., 2014).

How do we reconcile the increasing evidence that information about working memory contents appears in visual cortical areas, yet univariate responses do not uniformly emerge in these areas? In fact, here we show that mid-level visual cortical areas instead showed meaningful *decreases* in univariate activity with working memory load. Viewing these results in light of prior work, we hypothesized that the characteristics of the population level responses may alter whether an increase or decrease in mean population activity is observed. Specifically, we noted two key findings. First, prior work characterizing visual population receptive fields (pRFs) has shown that pRFs in different retinotopic ROIs have unique population-level characteristics. Specifically, Aqil et al. (2021) developed a modified normalization model to characterize population-level *suppression* and *compression*, and they found that early, relative to late, visual regions showed higher amounts of surround suppression and later, relative to early, visual regions showed higher amounts of compression. Second, we see hints in prior published work that these hypothesized differences in pRFs may sustain from encoding and into the working memory delay period. For example, in Sprague and colleagues’ (2014) work reconstructing the position of a remembered item, a negative surround is visible in the population-level model estimates for earlier cortical regions (V1-V4) but not for IPS.

Here, we hypothesized that changes to the surround suppression of pRFs in different cortical areas may yield differential effects on the net univariate response of a population as more memory items are added. We used an adapted normalization model from Aqil et al. (2021) to visualize what happens when memory load and surround suppression are manipulated in this normalization model (Figure 4A). The results of the model output are shown in Figure 4B as a function of the memory load (peaks for 1 or 4 items) and whether or not surround suppression is added to the normalization model. We found that without surround suppression added to population level pRFs, we see a steady increase in univariate activity (i.e., mean pixel intensity of the simulated image) as a function of memory load (Figure 4C); with surround suppression, this pattern of results “flips” and results in a net decrease to univariate activity with load. These simulation results highlight how a unified explanation when considering multivariate coding (i.e., competition among item representations within each ROI), could lead to opposing univariate signals measured globally from different ROIs.

**Figure 4.**
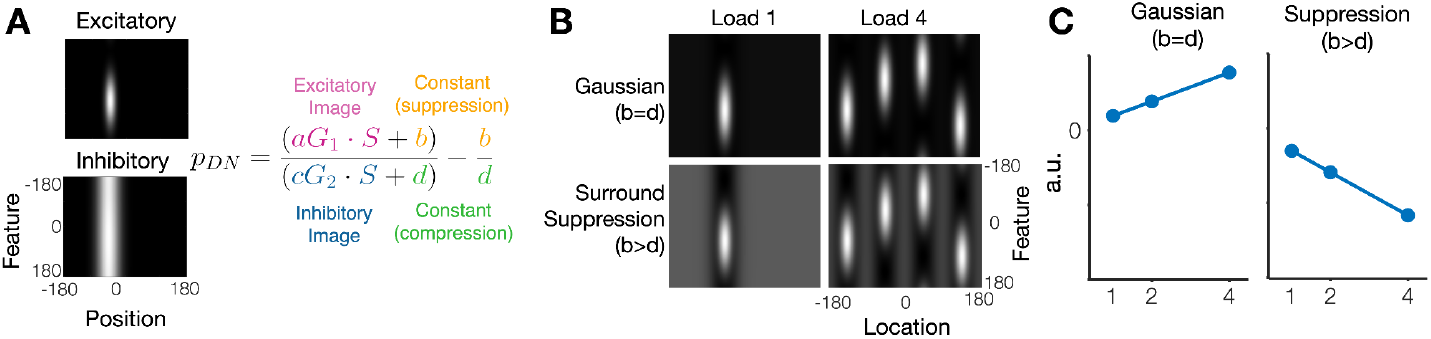
Simulating the effect of population surround suppression on univariate activity. (A) Adapted normalization model from Aqil et al. (2021). (B) Normalized images with a load of 1 versus 4 and with or without population level surround suppression. (C) Mean activity as a function of load, calculated as the mean luminance of pixels in each normalized image.

**Figure 5.**
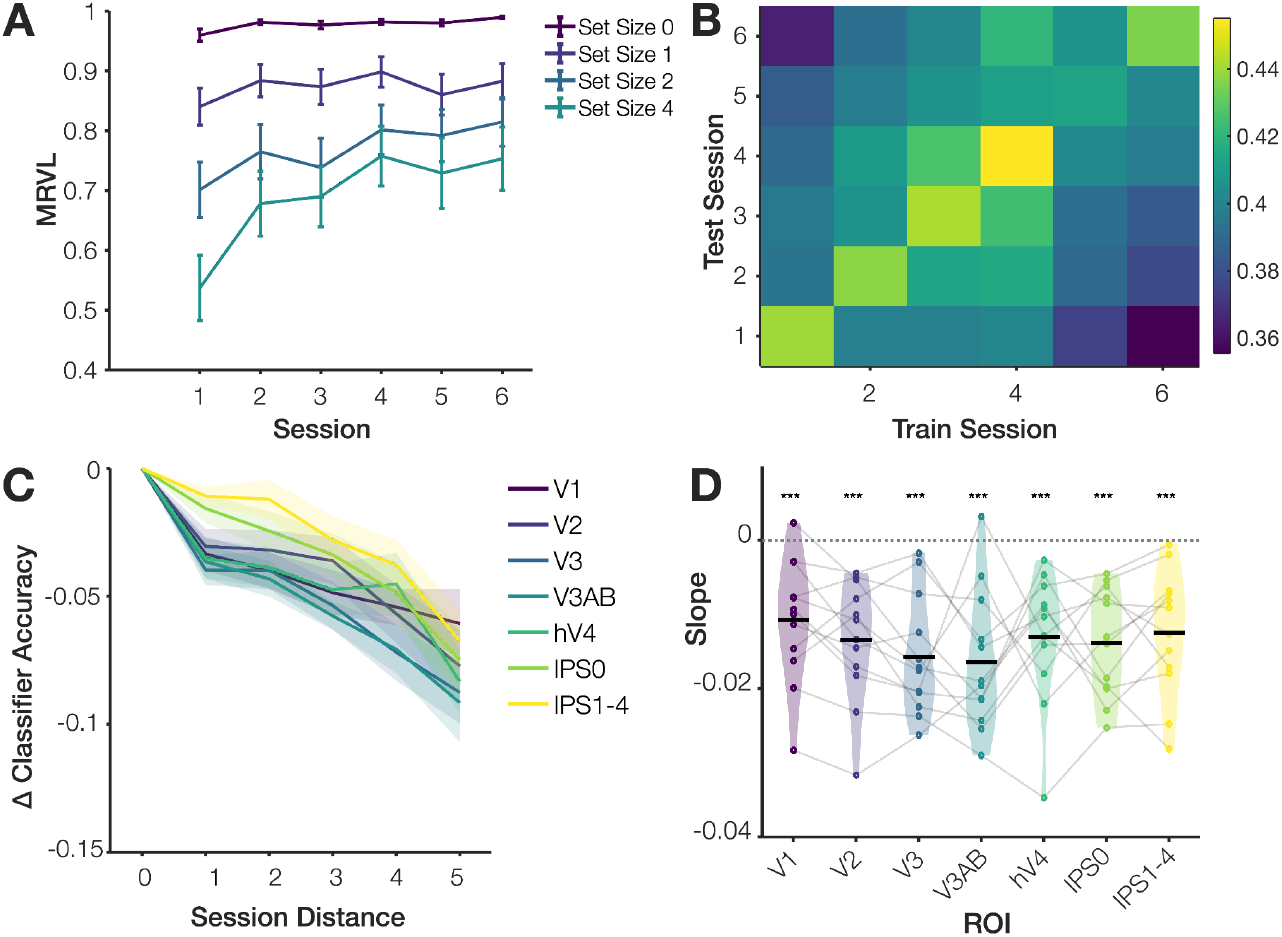
Cross-session generalization reveals representational drift of working memory load. (A) Behavioral performance (mean resultant vector length of error distribution) as a function of memory load condition and session number. Error bars represent 1 SEM. (B) Delay period load decoding, training and testing across sessions. (C) Decoding performance as a function of the distance between the training and testing sessions (0-5 sessions) for each ROI, baselined relative to a distance of 0 (within-session decoding). Shaded error bars represent 1 SEM. (D) Change in decoding performance as a function of (slope of lines in Figure 4C). Gray lines and dots represent individual subjects; violins depict the distribution; black lines represent the mean. *** p<.001

### Representational drift of multivariate load signals across weeks

The current dataset provided a unique opportunity to examine the possibility that multivariate prediction of working memory load may show directional drift over time.

#### Behavior over sessions

Participants’ performance significantly improved across the 6 sessions (Figure 1A). A repeated measures ANOVA with factors Load and Session revealed a main effect of Load, *F*(1.48,16.27) = 29.1, *p* < .001, *η^2^ _p_* = .73, a main effect of Session, *F*(5,55) = 13.1, *p* < .001, *η^2^ _p_* = .54, and an interaction between Load and Session *F*(15,165) = 6.0, *p* < .001, *η^2^ _p_* = .35. On average, participants completed 6 sessions over 79.4 days (SD = 50.1 days), with a mean inter-session interval of 27.5 days (SD = 19.4 days). The session timing for all participants is shown in Table S1.

### Generalizability of load decoding decreases monotonically as a function of the number of intervening sessions

To examine the generalizability of load decoding across seasons, we separately trained and tested classifiers on each pairwise combination of sessions (Figure 1B). Generalizability of decoding decreased as a function of the number of intervening sessions (Figure 1C); a repeated measures ANOVA revealed a main effect of Session Distance, *F*(2.61, 28.68) = 32.8, *p* < .001, *η^2^ _p_* = .75, a main effect of Region, *F*(2.74,30.19) = 46.2, *p* < .001, *η^2^ _p_*= .81,and no significant interaction, *F*(30,330) = 1.28, *p* = .15, *η^2^ p* = .10. For each ROI, we also calculated the slope, defined as the decrease in classification performance as a function of the number of intervening sessions between the training and testing data (Figure 1D). The average slope across ROIs was -.014 (SD = .0059), indicating a 1.4% decrease in decoding accuracy for each additional intervening session between training and testing.

### Supplemental Results

First, we expected to be underpowered to detect individual differences correlations, but we pre-registered that we would report these for completeness given previously reported correlations between univariate IPS activity and working memory behavior with similar sample sizes (Analysis S1). We did not observe any significant correlations between univariate activity and WM behavior after controlling for multiple comparisons. Second, we pre-registered that we would examine item-level responses using multivariate approaches used in past work during a 4-item visual search task (Adam & Serences, 2021); unfortunately, however, the current data did not support conducting these analyses as planned. Results from attempted versions of these analyses are shown in the supplemental results, and we further discuss potential issues and insights (Analysis S2).

## Discussion

In an fMRI experiment (n=12, t=12 scanner hours per person), human observers performed a working memory task while WM load was manipulated via a retro-cue (0, 1, 2 or 4 items). We examined univariate and multivariate signatures of working memory load in retinotopically-organized visual regions (V1 to IPS). We replicated landmark neuroimaging studies showing that the IPS shows sustained delay period activity that increases with working memory load (Todd & Marois, 2004; Xu & Chun, 2006). However, we also observed heterogeneous changes to univariate activity in other areas, including sustained univariate decreases to delay period activity as a function of memory load in areas V3, V3AB and hV4. Multivariate analyses revealed that, regardless of the univariate effect, information about the load condition could be decoded in all examined visual areas in a graded fashion (i.e., more similar load conditions were more confusable). In light of these results, we used simulations to illustrate how fundamentally converging multivariate models can, nonetheless, lead to diverging univariate effects with small changes to population-level parameters. Finally, we found that, across 6 sessions, multivariate information about WM load underwent directional drift, consistent with emerging findings of representational drift at multiple scales of analysis and for diverse aspects of behavior.

### Working memory load impacts univariate and multivariate signals across cortex

Although we replicated the prominent finding that univariate signals in the intraparietal sulcus (IPS) increase with working memory load, we did not find evidence to support the notion that the impact of working memory load preferentially impacts univariate signals in IPS. Instead, we found that information about working memory load could be decoded in areas from V1 through IPS regardless of the direction of the observed univariate effect in each region (increase, decrease, or null). Our observation of widespread load decoding converges with other emerging work suggesting that voxels responsive to WM load can be found distributed throughout cortex (van Ackooij et al., 2023) and with work demonstrating that working memory load can be distributed from coarse, scalp-level changes to the electroencephalogram (EEG) signal throughout the delay period (Adam et al., 2020; Jones et al., 2024; Suplica et al., 2025; Thyer et al., 2022). Similarly, our results fit with the broader finding from the sensory recruitment literature that the identity of a remembered item can be decoded from a distributed set of cortical regions from V1 through frontal cortex (Christophel et al., 2017; Ester et al., 2015; Serences et al., 2009). Together, our data suggest that the consequences of WM load are evident throughout the entire visual stream (i.e., starting from V1). We argue that load-dependent changes to univariate delay period activity are not diagnostic of a unique role for parietal cortex in generating working memory capacity limits. However, future work is needed to disentangle competing possibilities explaining the emergence of broadly-distributed load modulations in cortex. One possibility is that load modulations arise independently in disparate areas due to local processing constraints (i.e., competition and normalization among local populations of neurons)(Schneegans et al., 2020; Schneegans & Bays, 2017; Wei et al., 2012); however, it also may be that load modulations in earlier visual regions are inherited via top-down feedback originating in the fronto-parietal control network (Bouchacourt & Buschman, 2019; Panichello & Buschman, 2021; Park & Serences, 2025; Xu, 2023).

### Considering multivariate and univariate signals in tandem

Our simulations reveal the importance of considering potential interactive effects of multivariate and univariate signals in fMRI analyses. Using a simple simulation, we revealed how manipulating population response parameters (e.g., degree of surround suppression) can lead to superficial dissociations in univariate signals. For example, in much of the historic literature, a univariate increase versus decrease might be interpreted as opposing effects indicative of opposing underlying functions (i.e., usage goes up and this area and down in this other area, so these areas are oppositional in function or mechanism). However, our simulations suggest that opposing univariate signals can be created by changing model parameters but without implementing a fundamental change to the nature of the underlying model. Specifically, by manipulating the amount of surround suppression for population-level representations, we can “flip” the direction of the predicted univariate result as a function of memory load (from overall increase to decrease) without changing the core features of the underlying representation (e.g., normalization model with an activation peak representing the location and feature of each remembered item). This simulation reinforces the fraught nature of reverse inference (Poldrack, 2006), particularly for global univariate effects, and highlights the potential utility of considering both multivariate and univariate signals.

### Representational drift across sessions

Across our 6 sessions, we observed above-chance decoding of load condition when training and testing within each experimental session. Moreover, we observed a monotonic decline in classification accuracy as a function of the temporal offset between sessions. This monotonic decline indicates that poor across-session generalization does not simply reflect mean-reverting variability related to head motion, poor anatomical registration across days, or other nuisance factors. Instead, the continuous drop in across-session decoding accuracy is consistent with a steady drift in the activation patterns. Moreover, behavioral recall accuracy largely stabilized after the 2nd session for all set sizes, which does not rule out continual refinements in strategy, but does suggest that the initial rapid phase of learning was complete.

A great deal of recent empirical work demonstrates that neural codes also gradually drift over hours/days/months, even after measurable changes in behavioral performance have stabilized (Deitch et al., 2021; Driscoll et al., 2017; Kentros et al., 2004; Roth & Merriam, 2023; Schoonover et al., 2021). For example, single cells in mouse posterior parietal cortex encode the spatial position of the animal in an environment. However, the spatial selectivity of individual cells in the population changes dynamically over time such that a classifier trained to decode spatial position using data from one experimental session does not cross-generalize well to data collected in another experimental session (Driscoll et al., 2017). As in our data, cross-generalization decoding accuracy is not mean-reverting and instead drops monotonically as a function of temporal separation between experimental sessions, consistent with a directional movement of activation patterns in a high dimensional space (Driscoll et al., 2017; Schoonover et al., 2021).

Some theoretical work postulates that long-term representational drift reflects an adaptive process of searching the neural solution space for more accurate or sparse representations (Aitken et al., 2022; Kappel et al., 2015; Ratzon et al., 2024). For example, if task relevant information is initially encoded with a high degree of redundancy, then a slow process of sparsification via ‘drop-out’ might preserve just the most essential components that support task performance. Thus, even if behavioral performance does not measurably change, behavior might be supported by a more energy efficient and independent code that is less susceptible to interference from co-activated representations (Barlow, 1969; Ganguli & Sompolinsky, 2012; Levy & Baxter, 1996; Rolls & Treves, 1990; Wixted et al., 2014). Alternatively, others note that drift naturally arises in any high-dimensional system that exhibits learning or plasticity. For example, artificial neural networks learn by updating their weights via an algorithm such as stochastic gradient descent until the model’s output matches a target. In the high-dimensional weight space of such a model, a random walk will quickly move away from the origin with a very small probability of returning to the starting point (Domb, 1954). As a result, even undirected updates can produce non-mean reverting drift, and the system will continue to drift so long as the weight-updating rule is applied (Blanc et al., 2019; Ratzon et al., 2024; Yang et al., 2023). Indeed, some have proposed quantifying drift as a means to make inferences about the properties governing learning and plasticity in neural codes (Qin et al., 2023; Rule et al., 2020; Rule & O’Leary, 2022). Thus, non-mean reverting directionality is not sufficient to infer that drift plays a functionally meaningful role in shaping neural codes and further work will be required to link directional changes in neural codes with changes in processing efficiency or other functionally significant markers of information processing.

### Challenges to modeling individual item representations

In our preregistration, we had planned to quantify the representational fidelity of all individual items in the array using an inverted encoding model approach in order to test hypotheses about prioritization within working memory. We had previously used this multi-item modeling approach in prior work on visual search (Adam & Serences, 2021), but the approach did not work well for the current dataset. We think there are a few possibilities for this failure of the method to generalize. First, we expected more noise for a working memory task relative to a visual search task, given that the visual search task has a stronger sensory signal contributing to decoding. Although we offset this anticipated decrease to signal quality with additional fMRI sessions, the additional data acquired may have been insufficient to offset the increase in noise. Second, participants may not have maintained sufficiently strong color-space bindings. Although participants were required to report the color of each item at its corresponding location, they may have “recoded” the binding from a spatial code (i.e., red + upper left) to an alternate binding such as recoding from the precise spatial position to an abstracted codes (i.e., upper left quadrant = 1, then maintaining a verbal code of, “red + 1”). Based on prior work, we had assumed that participants would maintain strong spatial bindings even when the spatial information is not strictly task-relevant. For example, Foster and colleagues (2017) found that participants spontaneously maintained the spatial position, as decoded via EEG signatures of spatial attention, even when spatial position was entirely task irrelevant. However, other emerging evidence suggests that space may not be spontaneously maintained over the longer fMRI delay period (Thayer et al., 2025).

A third possibility is that the participants’ maintained a robust spatial code, but this spatial code was reformatted such that cross-generalization of decoding from the training task (1-item spatial WM task) to the test task (4-item retrocue WM task) was disrupted. Recent work has highlighted how orthogonalization of neural representations may be an effective means to reduce interference and to align representations with imminent behavioral goals (e.g., from perception to memory; with attentional selection; with task demands) (Iamshchinina et al., 2021; Kwak & Curtis, 2022; Li & Curtis, 2022; Panichello & Buschman, 2021; Xu, 2024). Thus, training a model across such transformations can lead to poor generalization (Iamshchinina et al., 2021). Although we trained and tested our model from one working memory task to another (1 item WM task → 4 item retrocue WM task), we think it is plausible that an analogous orthogonalization of codes may have occurred across our training and testing tasks. During the localizer task (training), participants perceived a single dot, and maintained the location of this dot across a blank delay. In contrast, during the main task (test), participants first encoded 4 colored squares into working memory, and then were cued to drop a subset of items. Thus, it may be plausible that difference in attentional prioritization and competition during encoding drove orthogonalization of model weights across tasks. Future work is needed to systematically test this possibility.

## Conclusion

Working memory is the central mental workspace for complex cognition, and its limited capacity means that even modest increases to working memory load (e.g., from 1 to 3 items) yield costs to behavioral performance. The neural and computational constraints underlying costs of working memory load have been a long-standing area of debate. Our results support a distributed and dynamic account of working memory load effects. First, we found evidence for distributed modulation of neural signals by working memory load, with areas across the visual stream showing hallmarks of working memory load. We replicated previous findings that the intraparietal sulcus (IPS) shows increased univariate activity with higher memory loads. However, we show that information about memory load can be decoded from all visual areas (V1 through IPS) regardless of whether individual regions showed increases, decreases, or no change in overall univariate activity. Using simulations, we then show that opposing univariate signals may be explained by changing the degree of surround suppression at the level of population codes, which can invert the pattern of univariate activity across increasing memory load. Second, we found evidence for dynamic and drifting memory load signals. Using cross-session decoding of memory load, we found that neural representations of memory load gradually drifted across experimental sessions, even after behavioral performance had largely stabilized. Together these findings show that working memory load effects are distributed throughout the visual processing stream - rather than localized to specific regions of parietal cortex - and are consistent with the hypothesis that population activity patterns are continuously refined over time to more efficiently support memory and behavior.

## Acknowledgements

We would like to thank Aaron Jacobson at the Keck Center for fMRI at UC San Diego for building the custom bore-mounted acrylic screen for stimulus presentation, and for his continual and generous support of researchers using the fMRI Center. We would also like to thank Tommy Sprague for sharing example Siemens Prisma multiband scanning protocols.

## Methods

In one pre-registered fMRI experiment, we measured working memory load signals in visual and parietal cortical regions. The full pre-registration document is available at: https://osf.io/h97jv Note, because we followed a pre-registration, some of the methods text is similar to or overlapping with the pre-registration text.

### Participants

We recruited a total of N=12 participants from the University of California San Diego and surrounding community (mean age = 27 years [SD = 2.3, min = 24, max = 31]; 9 female, 3 male; 10 right-handed) to participate in 6 fMRI sessions at the Keck Center for fMRI at UCSD. Each session was approximately two hours long, and participants were compensated $25/hour. Procedures were approved by the local IRB, and participants provided written informed consent.

### Stimuli and Task Procedures

Across 6, two-hour fMRI sessions, participants completed 50 runs of the main working memory task (800 trials total; 200 trials per set size), 20 runs of an independent, set size 1 working memory localizer (400 trials), and 8 runs of a sensory localizer (800 stimulus presentation events). Extra localizer runs were collected if time permitted. To identify retinotopically-defined regions of interest, participants also completed 1 additional 2-hour retinotopy scan (either as part of a prior study [N=10] or as a new retinotopy session [N=2]). Retinotopic mapping data were used to draw retinotopic regions of interest (V1-V3, V3AB, hV4, IPS0-4) following standard procedures (Engel et al., 1994; Swisher et al., 2007).

Stimuli were generated using MATLAB (2017b, The MathWorks, Natick MA) and the Psychophysics toolbox (Brainard, 1997; Kleiner et al., 2007; Pelli, 1997) on a laptop running Ubuntu 20.04. Stimuli were rear-projected onto a custom acrylic screen mounted inside the scanner bore and viewed through a mirror attached to the head-coil; the projected screen had dimensions of approximately 57 by 32 cm, with a viewing distance of ∼81 cm (19.9° to the left/right of fixation, 11.3° above/below fixation). Responses were collected with an MR-compatible gamepad controller from Current Designs, Inc (Philadelphia, PA).

#### Main working memory task

The main task was a retro-cue variant of a whole-report visual working memory task (Adam et al., 2017) in which participants encoded 4 items and then were cued to maintain either 0, 1, 2 or 4 items across a blank delay. After a blank inter-trial interval, participants encoded 4 colored squares (0.5 s). The squares were equidistantly placed around an imaginary circle around fixation (radius = 7°), with 1 square in each visual quadrant. After a brief inter-stimulus interval (0.5 s), a retro-cue indicated which of the encoded squares should be remembered (0.8 s). Each retro-cue display consisted of four small shapes placed on an imaginary circle around fiction (radius = 1.8°). The shape identity in each quadrant indicated whether the participant should prioritize, store, or drop the item at each position (e.g., circle = prioritize, diamond = store, triangle = drop). The relationship between shape identities and task instructions was counterbalanced across participants. Participants were instructed to remember all items with either a prioritize or store cue; the purpose of the “prioritize” cue was to ensure that the spatial location of participants’ first response would be somewhat spatially counterbalanced across trials. In previous datasets using free recall procedures, we have observed that participants showed a spatial bias to first report items in the upper left when they are free to recall items in the order of their choosing. After a blank delay period (11 s), participants recalled all remembered colors in the order of their choosing (4 s per response). For Load 1, 2, and 4 trials, participants reported the color of all stored items.For Load 0 trials (no squares were remembered), participants performed a perceptual matching control task, in which they clicked a color on the color wheel that matched a presented color. Participants also completed 1 perceptual matching response for Load 1 trials.

#### Independent sensory and spatial working memory tasks

In addition to the main working memory task of interest, participants also performed a sensory localizer task and a 1-item spatial working memory localizer task. In the sensory task, participants performed a fixation task while a flickering checkerboard appeared in the periphery. In the spatial working memory task, participants remembered the position of 1 dot across an 11 s delay. Additional information about these tasks is provided in the supplemental (Methods S1).

### fMRI Acquisition and Preprocessing

Participants completed 6 fMRI sessions (approximately 2 hours each), during which they completed functional runs of three different tasks. Data were collected using a Siemens Prisma 3T scanner at the Keck Center for Functional Magnetic Resonance Imaging on the UCSD campus. Functional echo-planar imaging (EPI) data were collected with a Siemens 32 channel head coil and the CMRR MultiBand Accelerated EPI Pulse Sequences. To acquire the functional images, we will use the following settings: Multiband (MB) 2D GE-EPI with MB factor of 4, 44 2.5-mm interleaved slices with no gap, voxel size 2.5mm, field-of-view (FoV) 200 × 200 mm, no in-plane acceleration, repetition time (TR) 750 ms, echo time (TE) 30 ms, flip angle 50 deg, posterior-anterior phase encoding, slice tilt determined by Siemens AutoAlign. Forward and reverse phase-encoding directions were used during the acquisition of two short “top-up” datasets. From these images, susceptibility-induced off-resonance fields were estimated (Andersson et al., 2003) and used to correct signal distortion inherent in EPI sequences, using FSL top-up (Jenkinson et al., 2012; Smith et al., 2004).

Pre-processing of imaging data closely followed published lab procedures using FreeSurfer and FSL (Adam & Serences, 2021; Rademaker et al., 2019). We performed cortical surface gray-white matter volumetric segmentation of a high-resolution anatomical volume (1 mm^3^ isotropic) from the retinotopy session using FreeSurfer’s “recon-all” procedures (Dale et al., 1999). The first volume of the first functional run from each scanning session was coregistered to this common T1-weighted anatomical image. To align data from all sessions to the same functional space for each participant, we created transformation matrices with FreeSurfer’s registration tools (Greve & Fischl, 2009), and we used these matrices to transform each four-dimensional functional volume using FSL’s FLIRT (Jenkinson et al., 2002; Jenkinson & Smith, 2001). After cross-session alignment, motion correction was performed using FSL’s McFLIRT (no spatial smoothing, 12 degrees of freedom). Voxelwise signal time-series were normalized via Z-scoring on a run-by-run basis.

We drew visual ROIs (V1, V2, V3, V3AB, hV4, IPS0, and IPS1-4) using the retinotopy session data following published lab procedures (Rademaker et al., 2019; Sprague & Serences, 2013). From these retinotopically-derived ROIs, we further analyzed the subset of voxels that were selectively active for the stimulus locations used in the spatial localizer task. To select voxels, we will ran a one-way ANOVA on each voxel with a factor for visual Quadrant on each voxel; significant voxels (p < .05 uncorrected) were retained for other analyses. Analyses were performed using MATLAB 2018b (The MathWorks) and Python 3.10.14. Key packages for Python analyses include Jupyter (Kluyver et al., 2016), pandas (McKinney, 2010), numpy (Harris et al., 2020), matplotlib (Hunter, May-June 2007), and sklearn (Buitinck et al., 2013).

### Analyses

#### Univariate load signals

We estimated the Hemodynamic Response Function (HRF) for each voxel to recover BOLD time courses for each load condition. For each retinotopic ROI, we quantified average delay period activity during the time window 5.25 to 12.75 s following memory array onset.

#### Multivariate load signals

We used a multinomial logistic regression approach (‘LogisticRegressionCV’ from sklearn) that facilitates hyperparameter tuning by cross-validation across regularization strengths (‘Cs’ from −10 to 0). We conducted this analysis within each time-point (moving window of 4 TR’s averaged), and we iterated the analysis using each run as held-out test data (i.e., on each iteration of the analysis, 49 runs were used as training data and 1 run was used as test data). Cross-validation for each unique training/test set was conducted with 6 folds, using the L-BFGS solver (solver=‘lbfgs’) for optimization under a ridge penalty (penalty=‘l2’). The analysis was parallelized across all available CPU cores (n_jobs=-1), and a maximum of 500 iterations (max_iter=500) was specified to ensure convergence. As pre-registered, we also first tested whether we could decode working memory load in each ROI using a simple linear classifier (‘classify.m’, ‘diagLinear’ covariance option, analysis conducted for each TR and iterated using each run as the held-out data). The two decoding approaches led to consistent conclusions (Figure 3; Figure S2).

#### Cross-session drift of multivariate load signals

We extended the multivariate load decoding analysis to examine changes to decoding over the 6 sessions. Specifically, we trained and tested the multinomial logistic regression classifier on each possible pair of sessions. The classifier was trained using N-1 runs from one session, and tested on 1 run from the other session. Most participants had 8 to 9 main runs each session across the 6 sessions. Note, one participant had one short session due to technical difficulty (3 runs) and a seventh make-up session. For this participant, we used the first 6 sessions for cross-decoding analyses, and we reduced the amount of training and testing data so that the amount of data would be consistent across all sessions (i.e., train on 2 runs, test on 1 run).

### Summary of deviations from pre-registration

#### Added

(1) We did not predict *a priori* that we would see load-dependent decreases in activity in visual cortex. To think through his result, we conducted simulations to better understand the potential relationship between univariate and multivariate population signals. (2) In light of emerging findings about representational drift across cortex during task learning in rodents (Bellafard et al., 2024; Driscoll et al., 2017; Marks & Goard, 2021), our multi-session dataset provides a unique opportunity to examine representational drift of working memory load signals in humans.

#### Altered

(1) One participant completed 7 sessions instead of 6 sessions (one session was interrupted; the participant completed the remainder of the session at a later date). (2) We originally planned to use aggregate region IPS0-3 for parietal cortex analyses. However, upon finishing our pre-registered analysis of individual regions, we saw that IPS0 did not contribute much to the observed IPS load effects. As such, we present individual regions (V1-IPS0) as well as an aggregate IPS1-4 region. (2) We originally planned on using a simple linear classifier to classify working memory load (‘classify.m’ function, ‘diagLinear’ covariance option). This analysis worked as expected to classify working memory load (see Figure S2). However, we decided to use a more robust multinomial logistic regression approach for the main decoding analyses (see Figure 3). The qualitative pattern of all results was the same regardless of classifier choice.

#### Dropped

(1) We originally planned to examine individual item representations using a multi-item inverted encoding model approach following Adam & Serences (Adam & Serences, 2021). However, there were unforeseen difficulties with this analysis that prevented interpretation of these analyses as planned. Please see the supplemental for a summary of the inverted encoding model analyses and why they were not informative for the planned hypotheses (Analysis S2; Figures S3 and S4).

## Footnotes

1 Greenhouse-Geisser corrected values are reported where the assumption of sphericity is violated (Mauchly’s test).

2 Note, Figure 3B is averaged across ROIs, because ROI did not interact with Load Condition Pair in an initial repeated measures ANOVA (p = .10).

