## Supplemental Information for "Distributed and drifting signals for working memory load in human cortex"

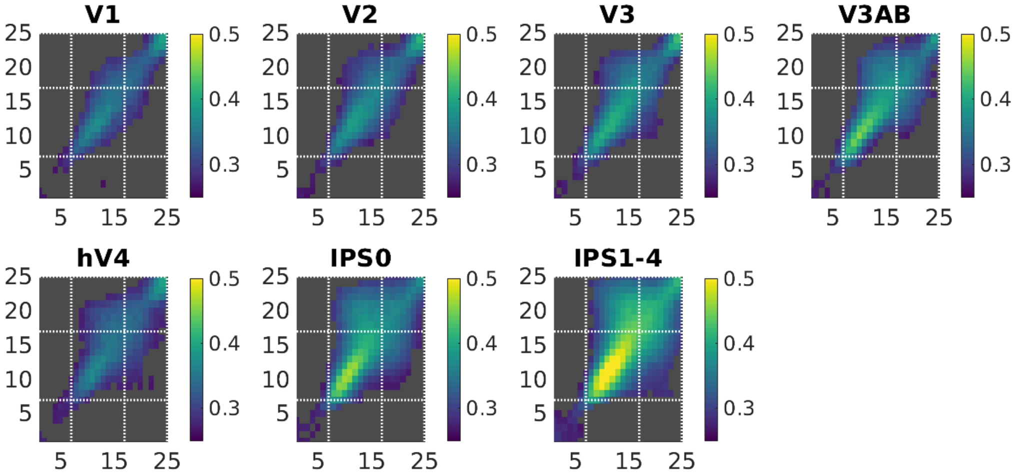


**Figure S1.** Cross-temporal generalization of load decoding (linear classify ('classify.m', 'diagLinear' covariance option; and decoding within each TR). Non-significant timepoints are masked in gray.

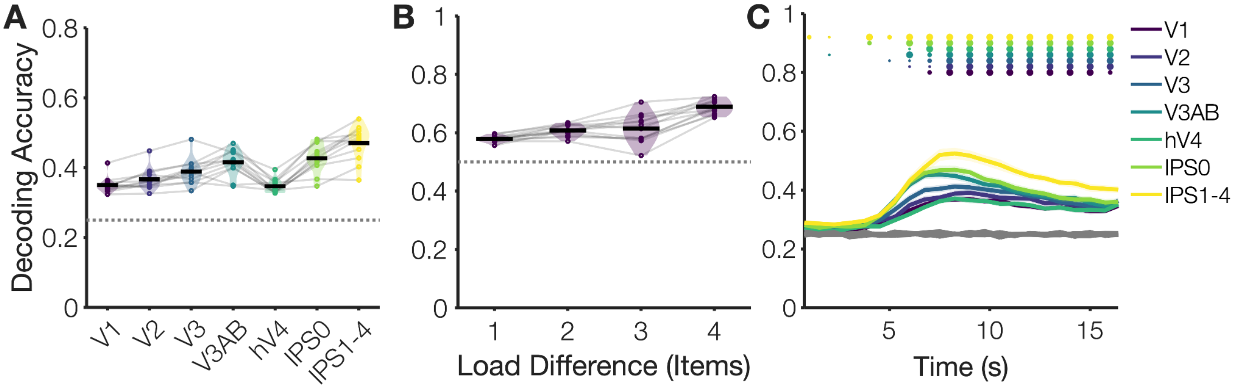


**Figure S2.** Load decoding with a linear classifier. Analyses from Figure 3 with a simple linear classify ('classify.m', 'diagLinear' covariance option), and decoding within each TR with no sliding window (every 0.75 ms). All key results from Figure 3 replicate including (A) above chance decoding in all regions (B) increase in decoding for pairs of conditions with a greater difference in memory load (e.g., 0 vs. 1 = difference of 1, 0 vs. 4 = difference of 4) and (C) sustained decoding throughout the delay period.

**Methods S1.** Additional methods information for the sensory and working memory localizers.

**Sensory localizer task.** Participants viewed a reversing checkerboard wedge (flicker = 4 Hz; white & black checkerboards) at one of 20 positions (equally spaced around a circle with radius = 7º). Each wedge's width along the circle filled 15° arc (width) and extended ~5° on either side of the imaginary circle (height). The flickering wedge stayed at each position for 3 s before moving to a new position (constraint: back-to-back positions must be in different quadrants). There were 100 wedge presentations within each run. Participants were instructed to attend to the fixation point; if the fixation point's color changed (increase or decrease in brightness), the participant pressed a button on the gamepad controller. A total of 20 fixation point color changes occurred throughout each run; changes to the fixation cross happened at random times with respect to wedge stimulus onsets.

**Working memory localizer task.** Participants remembered the spatial location of a single item (a gray dot) around the axis of an imaginary circle with radius = 7°. The position of the memory item could occupy any of 200 distinct values uniformly spaced between 0 and 360 degrees. To ensure a fairly even distribution of positions within each run, these 200 positions were binned into 20 bins of 10 positions each, and the 20 trials within each run were sampled from each bin. On each trial, the fixation point turned green to cue the participant that a memory item would appear soon (0.65 seconds). The memory item was shown for 0.5 seconds. After an 11 second delay period, the participant recalled the location of the dot by using buttons on the button box to move a "probe" dot around the circle until they think it best matches the location of the remembered dot (3 s response period; participants used the trigger buttons to move the probe quickly, and participants used the A and B buttons to move the probe slowly). The probe began at a random starting position on each trial, so that there was no relationship between the initial probe location and the true memory location. This ensured that the participant could not preplan a motor response). Each run of the spatial memory localizer included 20 trials with a jittered inter-trial interval (1-5 seconds) between each trial.

**Analysis S1.** Individual differences in univariate activity and working memory behavior.

With a sample size of 12 participants, we expected to be extremely under-powered to detect any potential brain-behavior correlations. However, given prior work reporting a relationship between IPS activity and WM behavior with similar sample sizes, N= 17 and N = 14, (Todd and Marois 2005), we pre-registered that we would report an individual differences analysis for completeness. We first calculated the correlation between average delay period activity for set size 4 and average working memory performance for set size 4. We did not observe any significant relationships after correcting for multiple comparisons [uncorrected p-values: V1 $r = -.09$, $p = .78$; V2 $r = -.20$, $p = .54$; V3 $r = -.20$, $p = .52$; V3AB $r = -.69$, $p = .01$; hV4 $r = -.08$, $p = .8$; IPS0 $r = -.45$, $p = .14$; IPS1-4 $r = -.45$, $p = .15$]. Furthermore, the direction of the correlation in IPS1-4 was opposite from what would be predicted based on prior work. A positive correlation indicates greater IPS activation corresponding with better working memory performance; a negative correlation indicates greater IPS activation corresponding with worse working memory performance. We also repeated this analysis when baselining the neural activity to set size 1 as has been done in earlier work (i.e., set size 4 delay period activity - set size 1 delay period activity). We observed one significant relationship after correcting for multiple comparisons [uncorrected p-values: V1 $r = -.11$, $p = .73$; V2 $r = -.54$, $p = .07$; V3 $r = -.51$, $p = .08$; V3AB $r = -.78$, $p = .003$; hV4 $r = -.51$, $p = .09$; IPS0 $r = -.28$, $p = .38$; IPS1-4 $r = -.03$, $p = .94$].

**Analysis S2.** Pre-registered analysis dropped from main paper: Decoding of item positions with an inverted encoding model **(Figures S3-S5).**

Following prior work (Adam & Serences, 2021; Brouwer & Heeger, 2009; Sprague & Serences, 2013) we used an inverted encoding model to estimate spatially-selective tuning functions from multivariate, voxel-wise activity within each ROI. We assume that each voxel's activity reflects the weighted sum of 20 spatially selective channels[1], each tuned for a different angular location. These information channels are assumed to reflect the activity of underlying neuronal populations tuned to each location. We modeled the response profile of each spatial channel as a half sinusoid raised to the $24^{th}$ power:

$$R = sin(0.5\theta)^{24},$$

where θ is angular location (0–359°, centered on each of the 20 bins from the mapping task), and  is the response of the spatial channel in arbitrary units.

Independent training data $B_1$ were used to estimate weights that approximate the relative contribution of the 20 spatial channels to the observed response at each voxel. Let $B_1$ (m voxels × $n_1$ observations) be the activity at each voxel for each measurement in the training set, $C_1$ (k channels × $n_1$ observations) be the predicted response of each spatial channel (determined by the basis functions) for each measurement, and W (m voxels × k channels) be a weight matrix that characterizes a linear mapping from "channel space" to "voxel space". The relationship between $B_1$, $C_1$, and W can be described by a general linear model:

$$B_1 = WC_1$$

We obtained the weight matrix through least-squares estimation

$$\widehat{W} = B_1 C_1^T (C_1 C_1^T)^{-1}$$

In the test stage, we inverted the model to transform the observed test data $B_2$ (m voxels × $n_2$ observations) into estimated channel responses, $C_2$ (k channels × $n_2$ observations), using the estimated weight matrix $\widehat{W}$ that we obtained in the training phase:

$$\widehat{C_2} = (\widehat{W}^T \widehat{W})^{-1} \widehat{W}^T B_2$$

[1] Note, in the pre-registration document we wrote that we would use 24 IEM channels. However, to attempt to optimize cross-decoding, we used 20 channels instead (such that the 4 stereotyped positions used in the main task will occupy the center of channels from the localizer tasks, i.e., 45, 135, 225, and 315 degrees)

Each estimated channel response function was then circularly shifted to a common center by aligning the estimated channel responses to the channel tuned for target location.

We trained the IEM using independent spatial memory task data (1 item) and test the model using single trial main task data (4 items). We then shifted and average the search task data so that like trials were aligned (e.g., "prioritized" item placed at position 0 within each set size condition). To reduce idiosyncrasies of only having 1 test set, we iterated the analysis by leaving out 1 block of training data and 1 block of test data, looping through all possible combinations (e.g., for each 1 block of left out training data, we left out each possible block of test data on different runs of the loop).

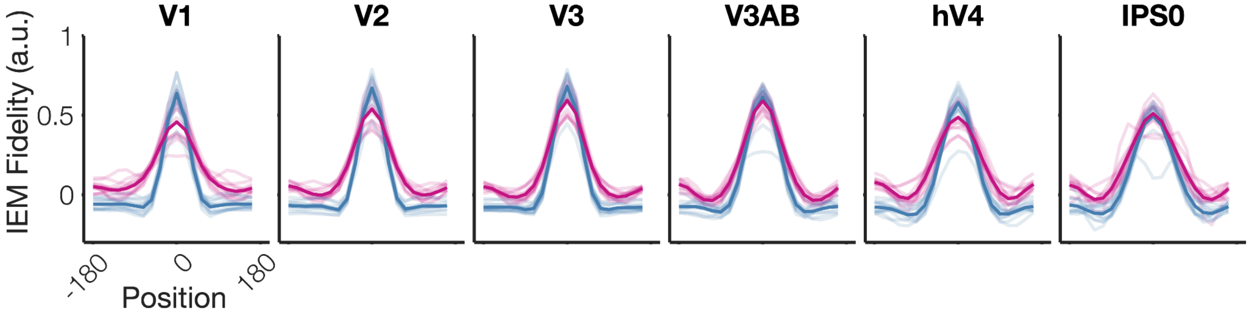


**Figure S3. Inverted encoding model results for localizer tasks.** Output of an inverted encoded model trained and tested within-task for each localizer. Blue: Trained and tested on sensory localizer task. Pink: Trained and tested on 1-item spatial working memory localizer task.

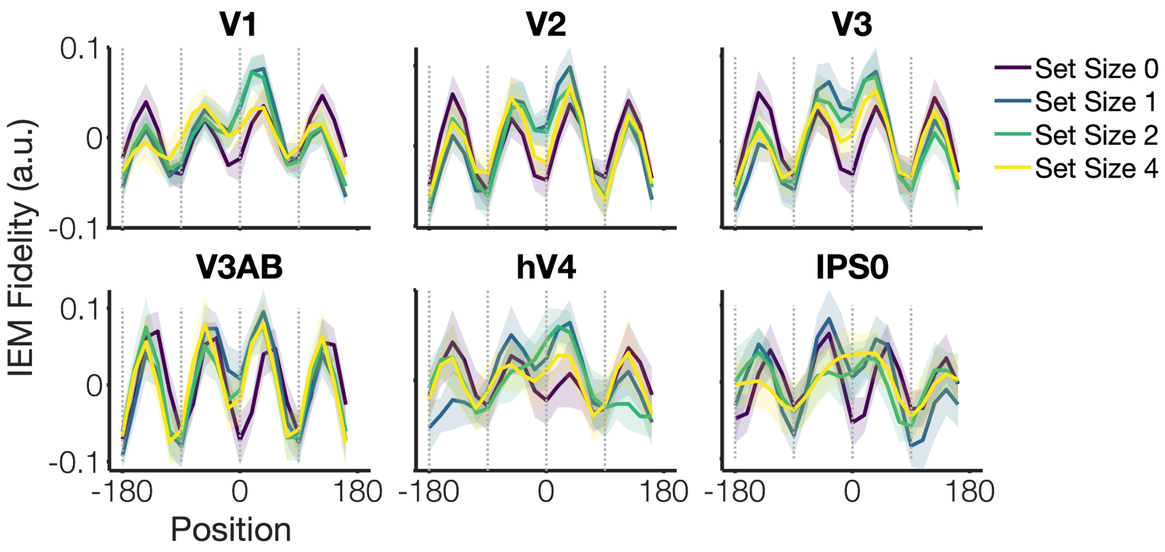


**Figure S4. Inverted encoding model results for main working memory task.** Output of an inverted encoding model trained on average delay period activity from a 1-item spatial working memory task and tested on average delay period activity from the main retrocue working memory task (4 items encoded, retrocued to maintain 0, 1, 2, or 4 items). Subplots indicate visual ROIs, and colored lines show model estimates for the 4 set size conditions (remember 0, 1, 2 or 4 items). Gray dotted lines indicate the expected “peaks” of remembered item positions at 0, 90, -90, and -180 degrees, where “0” represents the spatial position of the prioritized item. With strong generalization from the training task to the test task, we would have expected 4 “peaks” at the locations of the memory items (see: Adam & Serenes, 2021, *Journal of Neuroscience*). Instead, we observe “troughs” at the expected positions of items. This inversion may represent orthogonalization of spatial codes across the training/test tasks.

**Table S1. Days between sessions.** Time in days between each consecutive fMRI session for each participant.

| Subject | Session 2-1 | Session 3-2 | Session 4-3 | Session 5-4 | Session 6-5 | Session 7-6 | Session 6-1 |
|---|---|---|---|---|---|---|---|
| **1** | 12 | 8 | 113 | 29 | 4 | - | **154** |
| **2** | 12 | 2 | 6 | 8 | 21 | - | **37** |
| **3** | 14 | 83 | 15 | 19 | 35 | - | **152** |
| **4** | 6 | 23 | 14 | 21 | 5 | - | **63** |
| **5** | 5 | 8 | 13 | 16 | 8 | - | **45** |
| **6** | 5 | 16 | 27 | 56 | 62 | - | **161** |
| **7** | 8 | 7 | 6 | 18 | 8 | - | **39** |
| **8** | 7 | 21 | 14 | 19 | 9 | - | **63** |
| **9** | 5 | 40 | 20 | 15 | 35 | 7 | **110** |
| **10** | 7 | 14 | 7 | 17 | 3 | - | **41** |
| **11** | 7 | 8 | 5 | 30 | 5 | - | **48** |
| **12** | 11 | 21 | 7 | 7 | 5 | - | **40** |
| **MEAN** | **8.25** | **20.92** | **20.58** | **21.25** | **16.67** | - | **79.416667** |
| SD | 3.17 | 22.05 | 29.84 | 12.89 | 18.37 | - | 50.086214 |
